# Population dynamics of *Arabis alpina* in the French Alps: evidence for demographic compensation?

**DOI:** 10.1101/070847

**Authors:** Marco Andrello, Pierre De Villemereuil, Delphine Busson, Oscar Gaggiotti, Irène Till-Bottraud

## Abstract

- Due to its genetic proximity with *Arabidopsis thaliana, Arabis alpina* (Brassicaceae) is increasingly used as a perennial model species in studies of molecular evolution and adaptation.
- We studied the demography of *A. alpina* in six natural sites widely differing in their degree of disturbance, slope and vegetation, and encompassing the full altitudinal range of the species.
- We estimated three vital rates (growth, reproductive effort and survival) for individually-marked plants, studied for six years (2008-2014). We characterized the thermic conditions of each site with different thermal variables obtained using *in situ* continuous-time data loggers.
- Although *A. alpina* is described as a perennial species, the average life expectancy was only 1.82 years and most plants died before setting seeds. Plant size was a strong predictor of all three vital rates.
- Mean daily temperature showed a positive effect on growth and a negative effect on survival. Furthermore, reproductive effort covaried negatively with survival, suggesting a mechanism of demographic compensation acting on an elevational gradient.
- *Synthesis.* These results are informative of the selective pressures experienced by *A. alpina* in natural conditions and will help design experimental and molecular studies of local adaptation in this species.

## Introduction

The perennial herb *Arabis alpina* (L., Brassicaceae) is emerging as a new model organism in evolutionary developmental biology due to its proximity to *Arabidopsis thaliana* (Koch et al. 2006). *A. alpina* is distributed in the Northern hemisphere and lives at elevations going from few meters above sea level in Northern regions (Scandinavia) to several thousand meters in African mountains. In the European Alps, *A. alpina* is restricted between 800 and 3000 meters (Huxley 1986). Studies so far have focused on the genetic architecture of flowering and perenniality (Wang et al. 2009; Bergonzi et al. 2013), leaf senescence (Wingler et al. 2012; Wingler et al. 2015) and phenotypic traits as root hair and trichomes (Chopra et al. 2014). There is an ongoing effort in advancing the molecular knowledge of *A. alpina* by sequencing its genome (Melodelima and Lobreaux 2013; Lobreaux et al. 2014; Willing et al. 2015). The ecology of *A. alpina* has been addressed mostly from the perspective of molecular ecology through studies of its genetic structure and phylogeny (Assefa et al. 2007; Ansell et al. 2008; Ansell et al. 2011; Karl et al. 2012; Buehler et al. 2012) and detection of loci under selection (Poncet et al. 2010; Manel et al. 2010; Manel et al. 2012; Buehler et al. 2013; Zulliger et al. 2013; Buehler et al. 2014) or with an evolutionary ecology perspective aimed at characterizing its mating system (Tedder et al. 2015) and local adaptation (Toräng et al. 2015). Very few studies have tried to describe the population ecology of this species *in natura* and we lack basic knowledge on rates of survival and growth, reproductive effort and on the ecological factors driving the variation of vital rates in natural conditions.

In this study, we investigated the demography of *A. alpina* in six field sites across two distinct mountain ranges in the French Alps, encompassing the entire altitudinal range of the species. Environmental variables were measured *in situ* to reflect local conditions. Gaining knowledge and understanding on the population ecology of *A. alpina* will help scientists working on this species to formulate hypothesis about the selective factors that are likely to drive the evolution of the species in the wild. For example, studies aiming at identifying genes under selection are easier if the selective factors are known (Hancock et al. 2011). While laboratory studies can reveal some potential selective factors, field conditions are complex and selective factors may differ from those that become important in laboratory conditions (Bergelson and Roux 2010). This requires measuring vital rates (survival, reproductive effort and growth) *in natura,* describing their variation across individuals and geographical locations and understanding the ecological factors underlying such variations.

Vital rates of species vary in space and time in response to biotic and abiotic environmental conditions. Understanding how environmental conditions affect a species’ vital rates is one of the main topics in population ecology. This is however challenging because of the complex relationships between an organism and its abiotic and biotic environment. For well-studied model species (e.g. *Arabidopsis thaliana* and some economically important species such as crops), there is some knowledge on the eco-physiological processes that determine survival rates, fecundity and growth. For non-model species, this knowledge is often absent. In this case, a correlative approach can be employed: the correlation between some environmental variables and the vital rates of the species are tested and quantified. The choice of the variables is often done with very little a priori knowledge on the eco-physiology of the species. We took this approach to understand what environmental factors underlie the variation in vital rates (survival, fecundity and growth) in *A. alpina.* Specifically, we addressed three questions:

1. What are the demographic dynamics of natural populations of *A. alpina?*
2. What is the extent of variation in environmental conditions and population dynamics between different locations?
3. To what extent do environmental conditions affect population dynamics?

## Materials and Methods

### Study species

The artico-alpine plant *Arabis alpina* (L., Brassicaceae) occurs in most European mountain ranges, Anatolia, northern and eastern Africa and subarctic North America (Koch et al. 2006). *A. alpina* is often found in association with calcareous soil in disturbed habitats. In the European Alps, it is predominantly found in open and unstable grounds such as glacier forelands, scree slopes, rock ledges, and along foot paths and small streams in the montane, subalpine and alpine zones. It prefers moist sites with low vegetation cover. New seedlings germinate and establish throughout the growing season, plants flower for a few weeks and then flowers develop into fruits (siliques). The reproductive stems senesce and die while the plant overwinters as a rosette. Some degree of vernalisation is needed for the onset of floral development, which makes the plant perennial and iteroparous (Wang et al. 2009).

### Study sites

Six sites in three mountain ranges were studied (**Table 1**). Three sites (BRU, CHA and VIL) were located in the Vercors range, one site (LAU) was located in the Ecrins range near the Lautaret pass and two sites (GAL and PIC) were located in the Cerces range near the Galibier pass. For simplicity, we consider the Ecrins and Cerces sites as belonging to the same mountain range and refer to it as the Lautaret-Galibier mountain range. The maximal spatial distance between sites is 15 km in the Vercors and 4 km in the Lautaret-Galibier, while the average distance between mountain ranges is 65 km, thus supporting the choice of uniting LAU with GAL and PIC. The sites were located within a gradient encompassing the whole elevational range of *A. alpina* in the European Alps: the lowest site (BRU) was located at 930 m asl and the highest site (PIC) at 2930 m a.s.l. Besides elevation, the sites differed in aspect, soil type, slope, ground stability and vegetation (**Table 1**).

**Table 1.** Study sites

| Mountain range | Site name | Site code | # plots | Elevation<sup>a</sup> | Aspect | Latitude<sup>a</sup> | Longitude<sup>a</sup> | Soil type | Ground stability | Vegetation type |
| --- | --- | --- | --- | --- | --- | --- | --- | --- | --- | --- |
| Vercors | Gorges du Bruyant | BRU | 2 | 930 | S | 45.1534 | 5.6074 | Rock scree, calcareous | Unstable | Understory |
|  | Combe de Chaulange | CHA | 3 | 1480 | NW | 45.0693 | 5.5913 | Rock scree, calcareous | Stable | Shrubland |
|  | Villard de Lans | VIL | 3 | 1980 | SW | 45.0163 | 5.5700 | Rock scree, calcareous | Unstable | Subalpine prairie |
| Lautaret-Galibier | Lautaret | LAU | 3 | 2090 | N | 45.0268 | 6.3893 | Stream side, schistes | Very unstable | Under *Alnus viridis* cover |
|  | Galibier | GAL | 3 | 2590 | N | 45.0587 | 6.4028 | Shallow soil under rock cliff | Stable | Sparse |
|  | Pic Blanc du Galiber | PIC | 4 | 2930 | E | 45.0522 | 6.3838 | Rock scree, schistes | Very unstable | Very sparse |

<sup>a</sup>Latitude, longitude and elevation were measured with a GPS device. Elevation was further checked using the geographical coordinates on www.geoportail.gouv.fr.

### Demographic study

In each site, we set up two to four permanent quadrats to contain a total of roughly 50 individual *A. alpina* plants. The study started in 2007 in Vercors and in 2008 in Lautaret-Galibier, and ended in 2014. The position of each plant was marked with a small metal tag nailed into the ground and charted to facilitate the census. Once a year, during summer, we recorded for each plant the vital stage (alive or dead), the number of reproductive and vegetative stems and the number of siliques per reproductive stem. Stems bearing siliques or flowers were considered reproductive, while rosettes and stems bearing only leaves were considered vegetative. We attempted to census the sites at the time of fruit maturation, but because of differences in phenology among plants and among individual stems of the same plant, some stems would often bear flowers and some siliques would not be completely developed. We nonetheless included flowers and immature siliques in the fruit count. A sheep herd trampled the LAU site in 2010 killing the majority of the plants.

From census data, we computed the following variables for each individual and year *t:*

– *Plant size* (of year t). The sum of reproductive and vegetative stems of the plant in year t.
– *Growth* (from year t – 1 to year t). The ratio of number of stems in year *t* over number of stems in year t – 1.
– *Reproductive effort* (of year t). The total number of siliques of the plant in year t.
– *Survival* (from year t to year *t* + 1). A binary variable indicating survival from year *t* to year *t* + 1 (1 if the individual has survived and 0 if it has not).

To estimate seed set, we sampled 20 plants in each of three sites, VIL, LAU and GAL, outside but close to the permanent plots. We counted the seeds of each silique and averaged it over the siliques of the same plant (average seed set).

Plant age was determined only for individuals followed since their recruitment to the population, therefore from 2009 onwards. A power function describing the number of individuals as a function of age, *N* = *a.r^age^*, was fitted using nonlinear least squares in the R software. Life-expectancy was calculated as the average age at death across all cohorts.

### Thermal variables

We installed a temperature data logger (iButton^®^ hygrochron, Maxim Integrated™) in each quadrat at about 10 cm above ground and sheltered it from direct sunlight with a square and opaque Plexiglas plate. The data loggers recorded air temperature at the nearest half-degree every three hours. Data were recovered once per year during the annual census. Measures from quadrats of the same sites were highly correlated (R^2^ ≈ 0.90).

Some data loggers were lost and for some others the data was unreadable due to battery exhaustion or other unknown causes, especially in the last years of the survey. These data loggers were replaced with new ones; however, this resulted in loss of data. To replace missing data, we duplicated the data from the quadrats of the same site. When data were missing for all the quadrats of the site for a short period of time (23 occurrences), we reconstructed the missing data from nearby weather stations of the “Centre de Recherches sur les Ecosystèmes d’Altitude” (CREA – Mont-Blanc, creamontblanc.org): number 207 and 209 for the Vercors sites, and 218 and 29–217 for the Lautaret-Galibier sites. These stations were on average 12 km away from the closest study site (**Table S1**) and record temperature every 15 minutes. We fit a simple linear model with the temperature of the study site as response variable and the temperatures at ground level, 30 cm and 2 m above ground level of the two weather stations, as well as hour of the day, as explanatory variables. The models had a very good fit (average R^2^ = 0.82) and provided a reliable way to estimate the missing observations. To account for uncertainty in the estimates of the coefficients of the linear model, 10 replicated datasets were generated using values sampled from normal distributions defined by mean and standard error of the estimates of the coefficients

From the raw three-hourly measures, we calculated daily mean, minimal and maximal temperatures *(T_mean_, T_min_* and *T_max_).* These data were used to define thermal variables for the vegetative season; we disregarded the non-vegetative season (autumn and winter) because plants are dormant and less sensitive to environmental conditions. The limits of the vegetative season were derived from temperatures through the curve of the growing degree days, defined as *dd* = *(T_max_* + *T_min_)/2* - *T _base_*. We set *T_base_* = 0, because artico-alpine plants can grow at very low temperatures. For each quadrat, we determined graphically when the cumulative curve of *dd* started to increase and when it started to stabilize and used these two dates as the limits of the vegetative seasons (**Figure S1** in Supplementary information).

The vegetative season was further divided into two consecutive periods separated by the census date, to define thermal variables that had the same temporal structure as the census data. For simplicity, we refer to the first period as early season and the second period as late season. **Table S2** (Supplementary information) lists the starting and ending dates of the vegetative season and the census dates for each quadrat and year. Three-hourly and daily temperatures were used to define seven thermal variables for the early and late seasons (**Table 2**).

**Table 2.** Thermal variables

| Variables<sup>a</sup> | Description | Calculation |
| --- | --- | --- |
| T.min.1, T.min.2 | average daily minimal temperature | Mean of the daily minimal temperatures $T_{min}$ |
| *T.mean.1, T.mean.2<sup>b</sup>* | average daily mean temperature | Mean of the daily mean temperatures $T_{mean}$ |
| T.max.1, T.max.2 | average daily maximal temperature | Mean of the daily maximal temperatures $T_{max}$ |
| DD.1, DD.2 | accumulated growing degree days | Sum of the daily growing degree days $dd$ |
| *Frost.1, Frost.2* | number of frost days | Number of days with at least one three-hourly measure below zero |
| Frost.e.1, Frost.e.2 | number of frost events | A frost event is defined as a period of consecutive three-hourly measures below zero |
| *T.range.1, T.range.2* | average daily temperature range | Mean difference between daily maximal and minimal temperature, $T_{max} - T_{min}$ |

<sup>a</sup> Subscript 1 and 2 indicate early and late season, respectively.
<sup>b</sup> Variables in italics are those retained for the among-site comparison, the discriminant analysis and, except DD.2, as explanatory variables for the demographic variables.

### Statistical analysis

We analysed the thermal variables separately for the early and late season. Correlations between variables were measured by Pearson’s linear correlation coefficient. We reduced the number of thermal variables used in the subsequent analyses by retaining only the ones showing a pairwise Pearson’s correlation coefficient below 0.7 (**Table 2**).

Vital rates (growth, reproductive effort and survival) were analyzed using generalized linear mixed models including plant size and the uncorrelated thermal variables as potential explanatory variables. Two distinct sets of analyses were performed, a site-specific analysis and a global analysis.

In the site-specific analysis, we analyzed vital rates and environmental data separately for BRU, VIL and GAL (the other sites had too few observations to provide meaningful results). Quadrat and year were included as random effects. In the global analysis, we averaged vital rates, plant size and thermal variables over the years of the same site. Thus, we modeled the relationship between average thermal conditions and average vital rates. In the global analysis, we weighted the data points by the number of individuals used to compute the means to account for small sample sizes in some populations (e.g. PIC) and site was included as a random effect. In both the site-specific and the global analysis, the explanatory variables were standardized to facilitate the comparison of their effects.

Growth was modeled as a function of plant size in year *t* – 1, the environmental variables of the late season of year *t* – 1 and the environmental variables of the early season of year t. This choice of variables respect the time frame of the demographic process of growth, which takes place during the time from the census of year *t* – 1 until the census of year t. In the site-specific analyses, growth was log-transformed to achieve normality and variance homogeneity of the residuals, and plant size was log-transformed to obtain a log-linear relationship between plant size and growth, which corresponds to a power-function between size at time t-1 and size at time t. In both the site-specific and global analyses, the error structure was Gaussian.

Reproductive effort was modeled as a function of plant size in year *t* and the environmental variables of the early season of year t. In the site-specific analyses, the error structure was negative binomial to correctly model over-dispersed count data. In the global analysis, the error structure was Gaussian.

Survival was modeled as a function of plant size in year t, the environmental variables of the late season of year *t* and the early season of year *t* + 1, plus the interaction of all these variables with plant size. Ground stability was included as a categorical variable (**Table 1**) and the effect of sheep trampling in LAU in 2010 as a binary variable. In the site-specific analyses, the error structure was binomial. In the global analyses, the error structure was Gaussian.

Because of the uncertainty in the estimates of missing temperature measures by weather station measures, we used the 10 replicated datasets in a multiple imputation framework: we fit a model on each of the 10 dataset, averaged the estimates over models and computed the degrees of freedom according to Barnard and Rubin (1999). The efficiency of the multiple imputation framework was always above 0.99, indicating that 10 replicated datasets were largely sufficient to recover missing information. Because the multiple imputation framework relies heavily on t-tests, we selected the best models using a sequential t-test approach: the full model was fitted and variables with non-significant estimates were discarded; a new reference model was then fitted, and non-significant variables were again discarded until all remaining variables were significant. We then tried to add back all variables and created a new reference model with significant variables. These steps were repeated until a situation was reached were no variable could be added or removed to the reference model, which was then considered to be the best model.

All analyses were performed in R with the packages “ade4” v.1.7 (Dray and Dufour 2007), “lme4” v.1.1 (Bates et al. 2015), “mice” v.2.22 (van Buuren and Groothuis-Oudshoorn 2011) and “mitml” v.0.2.

### Demographic compensation

Demographic compensation was tested using linear models between two vital rates at a time (average survival, average growth, average reproductive effort) and weighting the data points by the number of individuals used to compute the means to account for small sample sizes in some populations.

## Results

### Thermal variables

On the basis of pairwise correlation coefficients among the initial seven thermal variables for each period (**Table S3 Supplementary information**), we retained only four variables for early season (T.mean.1, DD.1, Frost.1 and T.range.1) and three variables for late season (T.mean.2, Frost.2 and T.range.2) (Table 2).

**Figure 1** shows the variation in thermal variables among sites. Whereas variation among sites in the early season’s mean daily temperature was significant but small (T.mean.1, adj-*R*^2^ = 0.20, *F*_5,87_ = 5.5, *p* <0.001), the later season’s mean daily temperature clearly varied with site (T.mean.2, adj-*R^2^* = 0.82, *F*_5,67_ = 67.9, *p* <0.001) and elevation (T.mean.2 = *a* + *b* · Elevation, *a* = 12.77 ± 0.43 °C, *b* = 0.002 ± 0.0002, adj-*R*^2^ = 0.68, *p* <0.001).

**Figure 1.**
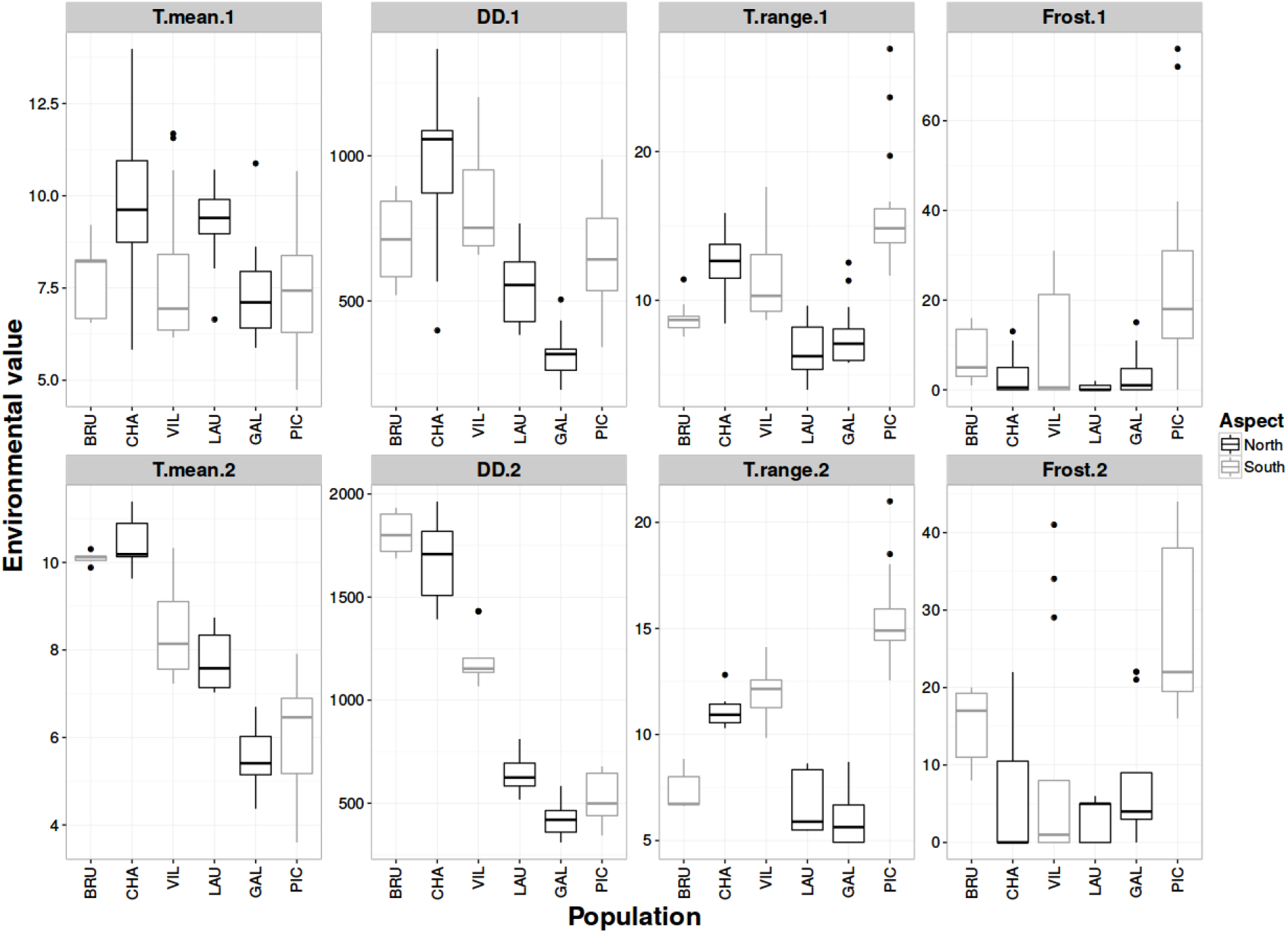
The four thermal variables in the six sites for early (top row) and late (bottom row) season.

The accumulated degree days were higher in the (lower elevation) Vercors sites than in the (higher elevation) Lautaret-Galibier sites, both in early (DD1 = 843 ± 234 dd in Vercors vs. 481 ± 229 dd in Lautaret-Galibier; adj-*R*^2^ = 0.37, *F*_1,95_ = 57.7, *p* <0.001) and late season (DD.2 = 1449 ± 289 dd in Vercors vs. 562 ± 130 dd in Lautaret-Galibier; adj-*R*^2^ = 0.82, *F*_1,71_ = 319.1, *p* <0.001).

The number of frost days was low on average, but PIC showed higher occurrences for the early season (Frost.1 = 24 ± 19 days in PIC vs. 5 ± 8 days in the other sites, *t* = 3.71, *p* <0.001) and especially for the late season frosts (Frost.2 = 34 ± 14 days in PIC vs. 9 ± 11 days in the other sites, *t* = 4.57, *p* <0.001).

Temperature range exhibited moderate differences between sites (adj-*R*^2^ = 0.60, *F*_5,87_ = 29, *p* <0.001 for the early season and adj-*R*^2^ = 0.87, *F*_5,67_ = 98, *p* <0.001 for the late season) and ranged from 9 ±1°C in BRU to 15 ± 3°C in PIC in the early season and from 6 ± 1°C in GAL to 16 ± 2°C in PIC in the late season.

PIC was thus the site experiencing the harsher conditions, i.e. large temperature ranges and high occurrence of frost events.

### Demography

VIL showed a decreasing trend in population size over 2008-2014 (**Figure 2**). In the other sites, population sizes decreased from 2008 to 2011, then increased until 2014. At the end of the study, population sizes of all sites except GAL were smaller than at the beginning. PIC nearly experienced extinction with only one individual remaining in 2012.

**Figure 2.**
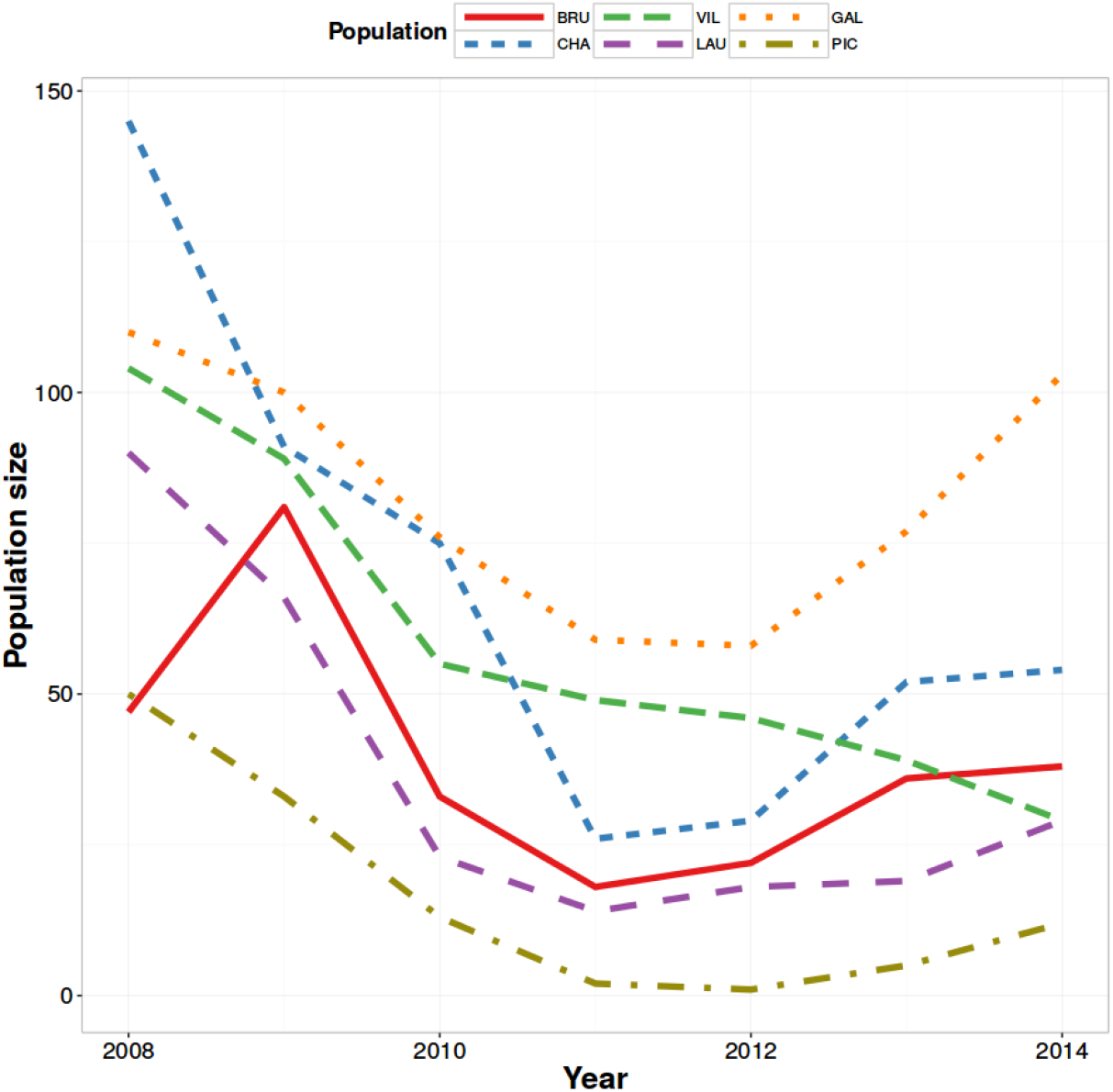
Number of *Arabis alpina* individuals in each site for each year of study.

The age distribution of plants was similar across sites, with decreasing number of individuals with age (**Figure 3**). The age distribution was well described by a power function *N* = a·r^age^, where *r* ranged between 0.23 (in PIC) and 0.53 (in GAL). Life-expectancy was 1.82 years (ranging from 1.42 in PIC to 2.09 in GAL). PIC was the site with shortest-lived plants, with 74% plants dying just after their first year, while the site with the longest-lived plants was GAL, where the plants dying just after their first year were only 48%.

**Figure 3.**
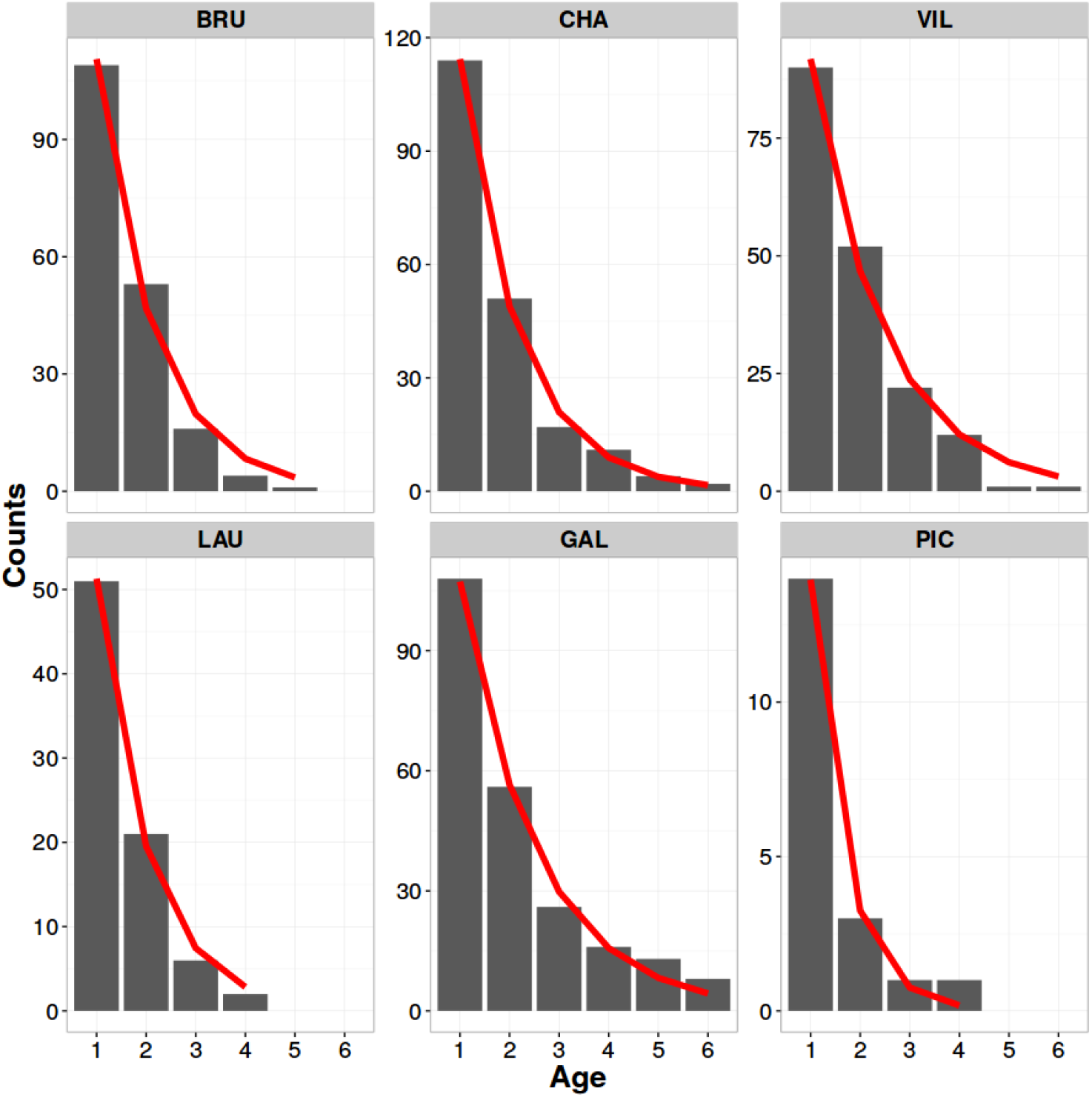
Number of *Arabis alpina* individuals in the six sites for each age-class, pooled over the six years of study. The red line is an exponential curve fitted to the data: *N* = *a.r*^age^

Average plant size ranged between 3.0 stems in LAU to 6.5 stems in CHA and was 4.4 stems on average. Most plants were small: the 75% percentile of the size distribution across sites was 5 stems. The average size of plants increased with age (**Figure 4**). However, in BRU and VIL the average size of older plants was smaller than the size of medium-aged plants and size reached a plateau in CHA.

**Figure 4.**
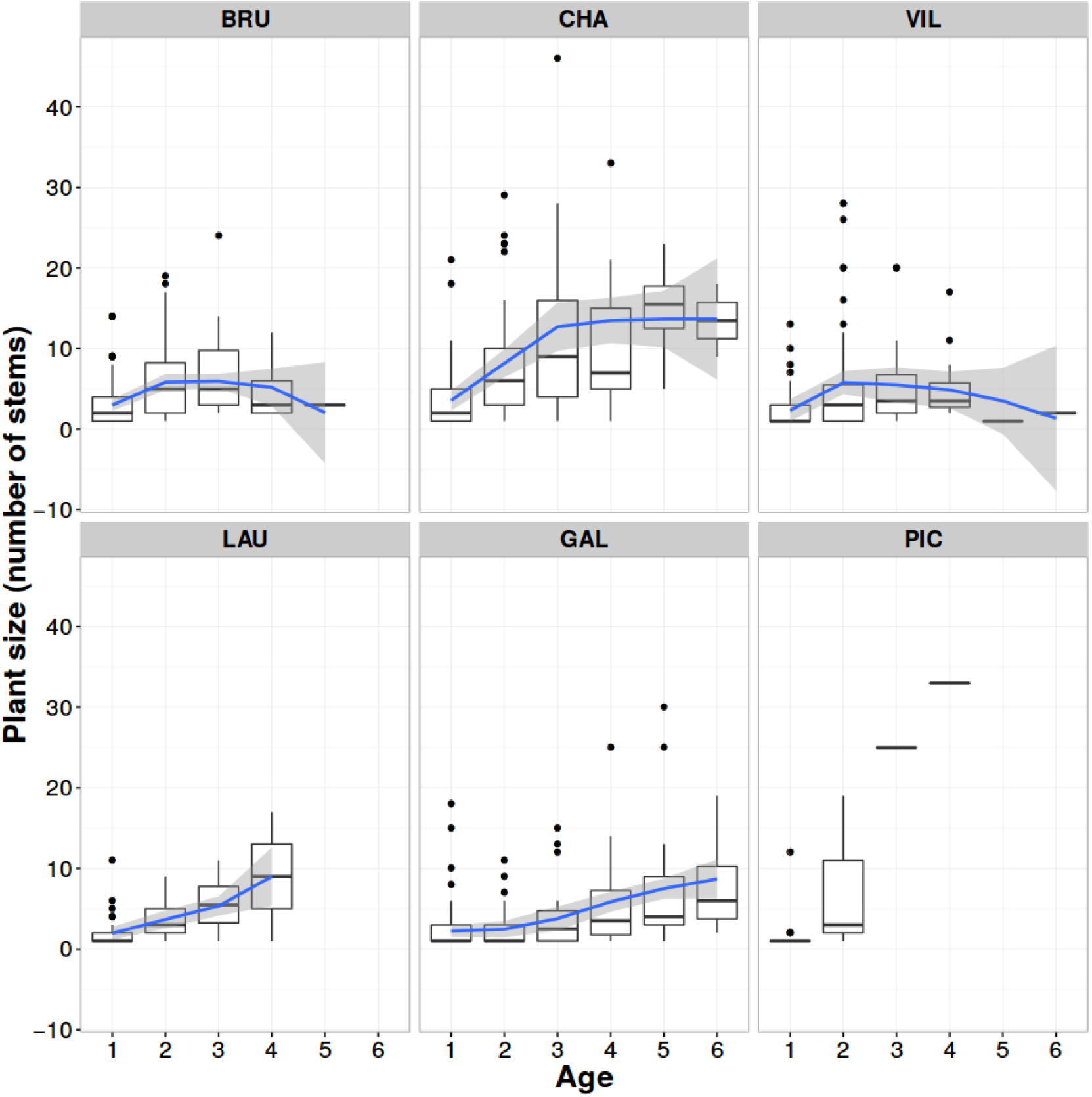
Distribution of plant size per site and age-class. The blue line is a local LOESS smoothing and the shaded grey areas are 95% confidence intervals of the LOESS algorithm.

Average survival rates increased with plant size, except in CHA, where the highest survival was observed for medium-sized plants. On average, 47% of plants died before becoming reproductive (ranging from 34% in BRU to 79% in PIC). Among reproductive plants, the average number of siliques per plant was 23.8 (ranging from 6.6 in GAL to 89.4 in PIC) and the average number of siliques per reproductive stem was 5.8 (ranging from 3.0 in GAL to 10.8 in PIC).

The average number of seeds per fruit was 25.1 and was not significantly different between the three studied sites (*F*_2,52_ = 0.036, *p* = 0.96).

### Site-specific analyses of vital rates

Growth was negatively dependent on plant size in the three sites (**Table 3**). Growth was positively associated to T.mean.1 in BRU and to DD.1 in VIL. These two variables were statistically linked in these two populations (Pearson’s linear correlation coefficient *r* = 0.85, *p* <0.001, in BRU; *r* = 0.67, *p* <0.001, in VIL), indicating a similar pattern of growth variation in these populations. Growth in GAL had a totally different behaviour with a strong negative association with Frost.1, as well as a positive association with Frost.2 and a negative association with T.range.2.

**Table 3.** Effects of plant size and thermal variables on temporal variation in growth of *A. alpina* for three sites (BRU, VIL and GAL)

|  | Estimate | SE | t value | df | p value |
| --- | --- | --- | --- | --- | --- |
| *BRU* |  |  |  |  |  |
| (Intercept) | 0.56 | 0.12 | 4.51 | 109.05 | < 0.001 |
| Log(Plant size) | -0.45 | 0.07 | -6.47 | 109.05 | < 0.001 |
| T.mean.1 | 0.35 | 0.07 | 4.77 | 109.05 | < 0.001 |
| *VIL* |  |  |  |  |  |
| (Intercept) | 0.16 | 0.09 | 1.81 | 140.04 | 0.036 |
| Log(Plant size) | -0.59 | 0.06 | -9.24 | 140.04 | < 0.001 |
| DD.1 | 0.14 | 0.08 | 1.84 | 140.04 | 0.034 |
| *GAL* |  |  |  |  |  |
| (Intercept) | 0.14 | 0.14 | 1.01 | 353.02 | 0.157 |
| Log(Plant size) | -0.27 | 0.03 | -7.96 | 352.73 | < 0.001 |
| Frost.2 | 0.09 | 0.04 | 2.35 | 350.72 | 0.010 |
| T.range.2 | -0.08 | 0.04 | -1.81 | 350.38 | 0.035 |
| Frost.1 | -0.17 | 0.04 | -4.76 | 351.43 | < 0.001 |

Reproductive effort in GAL was positively dependent on plant size and T.range.1 (**Table 4**). Models for reproductive effort in BRU and VIL did not converge.

**Table 4.** Effects of plant size and thermal variables on temporal variation in reproductive effort of *A. alpina* for GAL.

| GAL | Estimate | SE | t value | df | p value |
| --- | --- | --- | --- | --- | --- |
| (Intercept) | 0.19 | 0.30 | 0.65 | 451.01 | 0.257 |
| Plant size | 1.29 | 0.12 | 10.98 | 451.01 | < 0.001 |
| T.range.1 | 0.30 | 0.18 | 1.71 | 451.01 | 0.044 |

Survival showed the most complex pattern (**Table 5**). Notably, survival was positively correlated with plant size in VIL and GAL, but not in BRU. In fact, survival in BRU appeared to be only positively linked to one variable (T.mean.1). Inferences for VIL and GAL were more complicated with significant interactions with plant size, indicating that larger plants had a different sensitivity to thermal conditions than smaller plants.

**Table 5.** Effects of plant size and thermal variables on temporal variation in survival of *A. alpina* for three sites (BR, VIL and GAL)

|  | Estimate | SE | *t* value | df | *p* value |
| --- | --- | --- | --- | --- | --- |
| *BRU* |  |  |  |  |  |
| (Intercept) | -0.30 | 0.17 | -1.76 | 166.04 | 0.040 |
| T.mean.1 | 0.90 | 0.17 | 5.28 | 166.04 | < 0.001 |
| *VIL* |  |  |  |  |  |
| (Intercept) | -0.11 | 0.33 | -0.33 | 234.02 | 0.372 |
| Plant size | 0.54 | 0.17 | 3.16 | 234.02 | 0.001 |
| T.mean.1 | -0.54 | 0.17 | -3.19 | 234.01 | 0.001 |
| T.range.2 | 0.39 | 0.20 | 2.00 | 233.95 | 0.024 |
| Plant size : T.mean.1 | 0.45 | 0.19 | 2.37 | 234.02 | 0.009 |
| *GAL* |  |  |  |  |  |
| (Intercept) | 1.39 | 0.15 | 9.31 | 355.92 | < 0.001 |
| Plant size | 0.46 | 0.18 | 2.54 | 355.87 | 0.006 |
| T.range.2 | -0.53 | 0.13 | -4.00 | 355.58 | < 0.001 |
| DD.1 | 0.26 | 0.14 | 1.84 | 354.01 | 0.033 |
| Plant size : Frost.1 | 0.47 | 0.21 | 2.25 | 355.84 | 0.012 |

### Global analyses of vital rates

Average growth depended strongly and negatively on plant size, indicating that relative growth decreased as plant size increased (**Table 6**). Average growth depended positively on T.mean.2 and negatively on T.range.2 with comparable effect sizes: indeed, growth was much larger in the warmer populations BRU and CHA, and smaller in the colder populations GAL and PIC. Notably, growth depended only on late-season variables of the previous year, while early season variables of the current year were not significant.

**Table 6.**
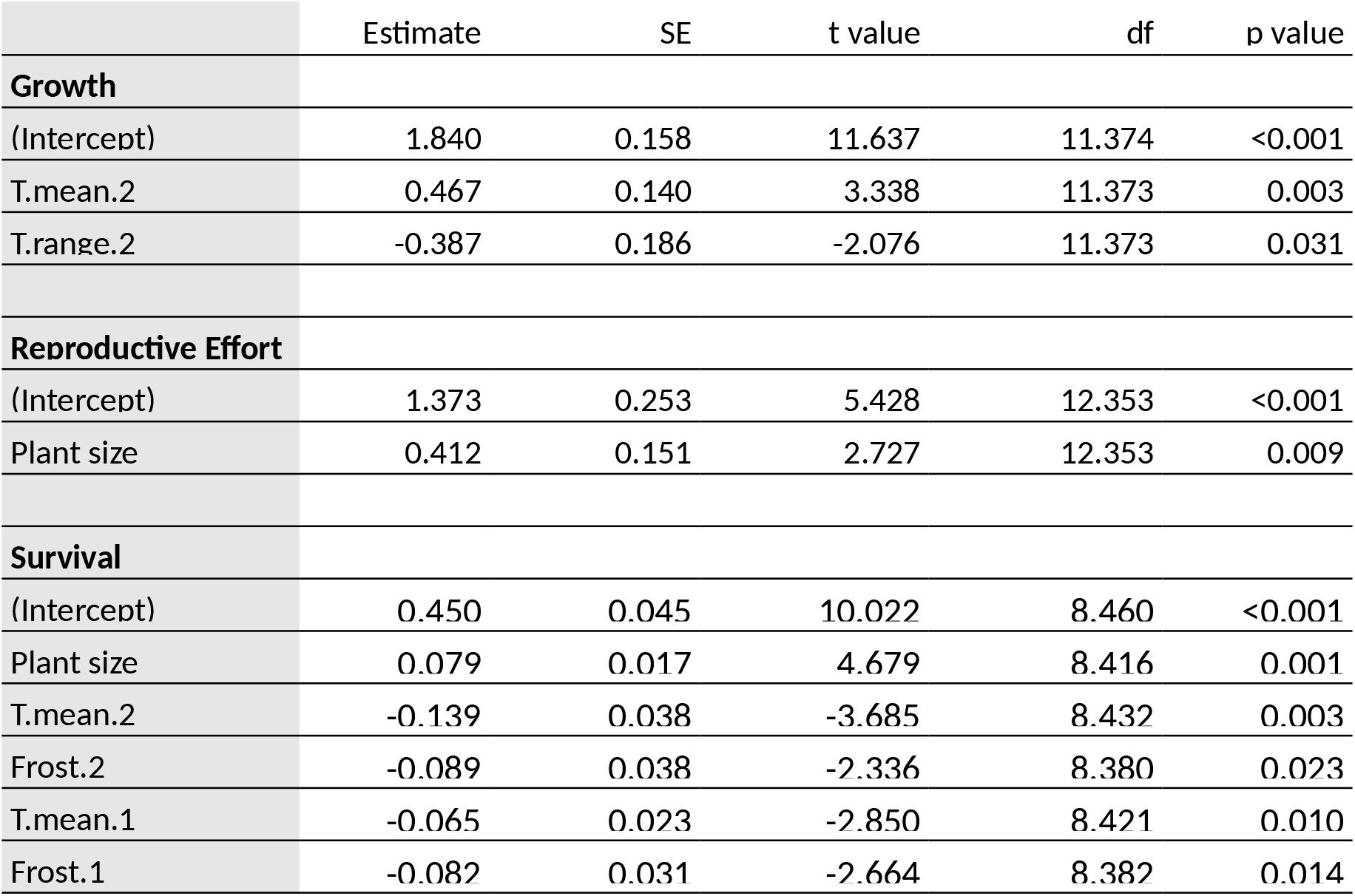
Effects of plant size and thermal variables on spatial variation in average vital rates (growth, reproductive effort and survival) of *A. alpina*

Average reproductive effort depended negatively on plant size, all other effects being non-significant. GAL was the only site with consistently lower reproductive effort compared to the other sites (ANOVA between sites, *F*_5,12_ = 3.32, *p* = 0.041; GAL effect, *t* = -3.49, *p* = 0.0045).

Average survival was positively affected by plant size and negatively affected by T.mean.1, T.mean.2, Frost.1 and Frost.2. Average daily mean temperature in late season (T.mean.2) had the strongest effect. Indeed, this variable is an excellent predictor of average survival, when excluding LAU and PIC (*R^2^* = 0.82), whereas the other variables allowed to account for the particular pattern of LAU (T.mean.1) and PIC (Frost.1 and Frost.2).

### Demographic compensation

Average survival was negatively, but not significantly, correlated to average growth (a = 0.75, *b* = -0.11 ± 0.07, adj-*R*^2^ = 0.08, *F*_1,16_ = 2.41, *p =* 0.14, **Figure 5**). The correlation was slightly stronger when LAU and PIC were removed from the analysis *(a* = 0.81, *b* = -0.13 ± 0.08, adj-*R*^2^ = 0.13, *F*_1,12_ = 2.97, *p =* 0.11, see Figure 5).

**Figure 5.**
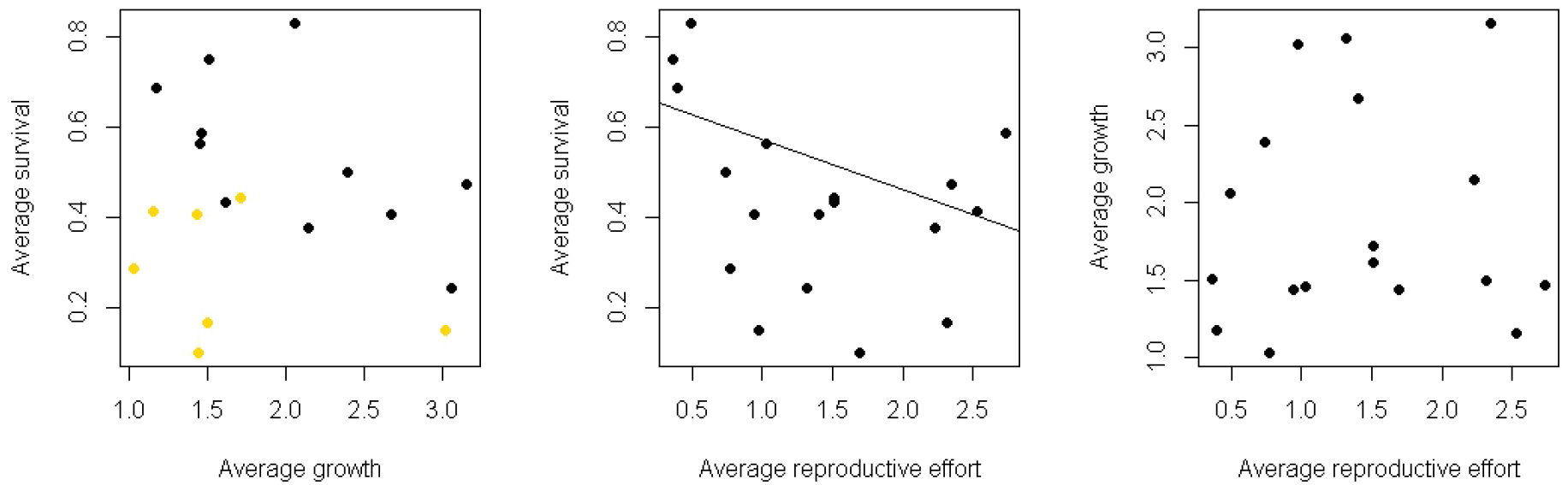
Pairwise correlations between average survival, average growth and average reproductive effort of *A. alpina*. Yellow points in the left panel corresponds to the LAU and PIC sites. The line in the middle panel is a fitted linear weighted regression.

The relationship between average survival and average reproductive effort was negative and significant *(a* = 0.68, *b* = -0.11 ± 0.05, adj-*R*^2^ = 0.18, *F*_1,16_ = 4.66, *p =* 0.04). However, the pattern was not linear and better described as a U-shaped (Figure 5).

The relationship between average growth and average reproductive effort was not significant (adj-*R*^2^ = 0.01, *F*_1,16_ = 1.2, *p =* 0.29).

## Discussion

### Demography

The six natural sites in which we have studied the demography of *Arabis alpina* differed strongly in their abiotic conditions. Slope, aspect, ground stability and thermal conditions varied greatly among sites. Despite these differences, the six sites showed some similarities in demography, particularly in the temporal variation of plant number during our study (**Figure 3**).

Although *A. alpina* is described as a perennial (Wang et al. 2009; Bergonzi et al. 2013; Wingler et al. 2015), life expectancy was only 1.8 years on average. We observed variation in life-expectancy, with some plants persisting for the entire duration of our study (i.e. up to 6 years), but at least half of the plants (up to 74% in PIC) died in their first year, with the consequence that half of the plants (up to 79% in PIC) never set seed. This high mortality is not an artefact of high seedling mortality, which is expected to be naturally high, because plants were marked only when they had several developed leaves. Life-history theory predicts that age at first reproduction decreases as mortality increases (Roff 2001), therefore earlier reproducers should be favoured in sites as PIC where mortality is high. In many alpine species, the timing of reproduction is regulated by vernalisation requirements and the genes controlling the plant responses to vernalisation have been studied in detail in *A. alpina* (Wang et al. 2009; Bergonzi et al. 2013). If functional genetic variation exists at these genes, alleles relaxing vernalisation requirement or conferring completely annual habits may be favoured in sites characterized by high plant mortalities. Yet another selective pressure would be the increase in resources limitation with altitude, implying a life-history response in which growth and reproduction are slowed down and adult survival and lifespan is increased (Körner 2003), as is apparently the case in the population GAL. As a consequence, the fitness advantage of an annual mutant should be evaluated considering all the components of the life-cycle and in particular the advantages of growing larger to produce more seeds, and the increasing survival rates at larger sizes (Horvitz and Schemske 2002). Indeed, for some years in PIC only a single large individual, bearing hundreds of siliques, was reproductive. More information regarding adaptive characteristics (e.g. fitness and genetics of life-history traits) in PIC population would be required to disentangle both evolutionary scenarios.

Average plant size increased with age. In some sites, average plant size showed a decreasing or stabilising pattern at older ages. This was the result of year-to-year variation in the number of stems of individual plants combined with a small sample size of long-living plants. Indeed, individual plant size in a given year was the sum of the number of stems developed in the same year and the number of stems overwintering from the previous year. Reproductive stems always die at the end of the reproductive season and cannot overwinter, while vegetative stems can overwinter but are subject to winter stochastic mortality. For many individuals, the obligate mortality of reproductive stems very often led to a decrease in plant size, because the new stems were not sufficient to replace the old ones. As a result, size variation was widespread in our data, which combined to issues of small sample size of long-living plants, sometimes caused this negative trends (in BRU and VIL, this trend is caused by <10 individuals).

We defined plant size as the number of stems without considering variation in biomass among stems, which was important but would have been too time-consuming and invasive to measure. Thus, patterns of variation in plant size should not be interpreted as a proxy for biomass variation or fitness. Nonetheless, the number of stems was useful to explain variation in survival, growth and reproductive effort. We thus advise using the number of stem as a simple and cost-effective metric to predict individual plant demographic rates of *A. alpina,* but we also encourage further studies of alternative metrics that may be better correlated to vital rates (Caswell 2001), such as plant diameter or the sum of stems weighted by the number of leaves.

Reproductive effort, measured as the number of siliques per plant, was highly variable and depended on plant size. Large plants could bear hundreds of siliques. The number of siliques per stem was less variable and ranged between 3 and 10. Each fruit bore on average 25 seeds and the number of seeds per fruit showed larger variation between individuals than between sites (only three sites studied).

### Effect of thermal variables on vital rates

Explaining variation in plant growth, survival and reproduction on the basis of topographical and geographical site characteristics is challenging because plant life is the result of many interacting factors such as elevation, aspect and microtopography (Körner 2007). This is why we decided to use local environmental data collected with data loggers located at vegetative height (a few cm above ground). This avoids the confusion of using environmental data from weather stations located many km away, which may not be representative of the environmental conditions experienced by the plants. Nonetheless, when looking at short periods of time, temperatures measured in nearby weather stations were correlated to temperatures measured *in situ* and could be used to reconstruct missing data within the vegetative season.

We used thermal variables estimated on periods that are relevant for the vegetative cycle of the plants rather than relying on calendar-based measures. Studies on alpine plants often use calendar-based variables such as monthly mean temperatures or annual minimal temperature to infer biogeographic patterns of species occurrence (Austin and Van Niel 2011) or local adaptation (Manel et al. 2012). While these data may lead to significant correlations at large spatial scale, it is difficult to interpret them at the scale of individual sites because the same calendar month corresponds to different phenological phases in different sites. Our choice of defining the vegetative phase on the basis of *in situ* temperature instead of calendar dates proved more realistic.

Using these data, we studied the relationships between vital rates (growth, reproductive effort and survival) and thermal variables. The effects of thermal variables on temporal variation in vital rates were not consistent across sites. We found comparable results for only two sites and one vital rate (growth rate for BRU and VIL, both located in the Vercors mountain range). Relationships between survival and thermal variables were highly site-specific. As survival is an integrative trait (i.e. influenced by many features of the individuals) and strongly submitted to local environmental and stochastic factors (e.g. random death due to accident, predation or disease), it might be expected that its variations are to be highly influenced by the local conditions.

Spatial variations in vital rates exhibited a clearer pattern. Spatial variation in average growth and survival rates were dominated by the mean daily temperatures of late season (T.mean.2), which strongly covaried with elevation (Pearson’s linear correlation coefficient *r* = -0.90, p <0.001). Growth rate was further affected by late season’s mean daily temperature range (T.range.2). This indicates that early environmental conditions have little effect on plant growth and, therefore, that growth mostly happens during the late season. However, this relationship may also be the result of smaller between-site variation in early season than in late season thermal variables (**Figure 1**). In addition to T.mean.2, average survival was affected by early season’s mean daily temperature (T.mean.1) and by the number of frost days in the early and late season (Frost.1 and Frost.2); a refined analysis showed that these latter variables explain average survival for the two sites not following the T.mean.2 trend (LAU and PIC).

More specifically, the relationship between growth and T.mean.2 was positive, whereas the relationship between survival and T.mean.2 was negative. A negative relationship between survival and growth would imply demographic compensation (Doak and Morris 2010, Villelas et al. 2015). Demographic compensation is implicated in shaping species distribution limits, where decreases in one vital rate at the species edge are compensated by increases in another vital rate. Over an altitudinal range or an environmental gradient, the mechanisms should be similar. We observed a negative relationship between survival and growth, but it was not significant, because of the small number of sites in our study. However, should this pattern be general, it would mean that *A. alpina,* at the specific level, follows a known rule of temperate alpine ecology, namely that high elevation plants tend to have higher survivals and lower growth rates than lower elevation plants (Körner 2003), because of physiological (e.g. lower temperature, lower partial CO_2_ pressure) and ecological (e.g. shorter reproductive seasons) processes. We observed however a significant negative relationship between survival and reproductive effort. In this context, the case of the GAL site is interesting. Despite not being the highest site, it displays strong signs of the syndrome described above, since it had the highest survival rate and the second lowest growth rate and the lowest reproductive effort. Moreover, GAL was the only site showing a lower average reproductive effort than the other sites, and reproductive effort was not related to thermal variables.

Despite similar temporal patterns of population dynamics among sites during the study (Fig. 3), vital rates had a highly site-specific behavior and the relationships between vital rates and thermal variables do not appear to be fully generalizable across sites. However, an effect of the late season’s mean daily temperature seems to emerge from the analysis of spatial variation in average vital rates (at least for growth and survival), although site-specific behavior again hampers strong conclusions regarding this phenomenon (especially for survival). More sites would be needed to ensure the generality of the observed trade-off between growth and survival. Yet, the possible existence of this adaptive compromise already raises the issue of its evolutionary origin, which could be either trough phenotypic plasticity or local adaptation. This would justify at the phenotypic level the numerous evolutionary genetic studies performed on this alpine plant (Poncet et al. 2010; Manel et al. 2010; Buehler et al. 2013; Buehler et al. 2014).

## Acknowledgements

We wish to thank Serge Aubert, Rolland Douzet and Bénédicte Poncet for helping us find the populations, Philippe Choler for interesting suggestions on the thermal determination of the vegetative season, Anne Delestrade for providing us with the weather station data and all the people helping with the field work, in particular Stefan Willhoit, Blaise Tymen, Elsa Jullien, Jérémy Camazzola, Anne-Lise Bartalucci, Lucas Hemery, Julie Chauvin, Florian Alberto, Loïc Chalmandrier, Aymeric Pilleux, Perrine Augrit and Justine Bisson. MA and PdV were funded by a PhD scholarship by the French Ministry of Research. This research was partially conducted at the Station Alpine J. Fourier, a member of the AnaEE network (Infrastructure for Analysis and Experimentation on Ecosystems).

## Supplementary information

**Table S1.** Location of the weather stations used to impute temperature data.

| Station ID | Latitude | Longitude | Mountain range | Distance to the closest site |
| --- | --- | --- | --- | --- |
| 207 | 45.12973N | 5.58719E | Vercors | 3 km |
| 209 | 45.02010N | 5.61786E | Vercors | 4 km |
| 218 | 45.02015N | 6.63496E | Lautaret-Galibier | 19 km |
| 29-217 | 44.83946N | 6.49312E | Lautaret-Galibier | 22 km |

**Table S2.** Dates of beginning and end of the vegetative season and dates of census.

| Site | Quadrat | Year | Beginning of vegetative season | Census date | End of vegetative season |
| --- | --- | --- | --- | --- | --- |
| BRU | 1 | 2009 | 31 March 2009 | 18 June 2009 | 01 December 2009 |
| BRU | 1 | 2010 | 21 March 2010 | 07 June 2010 | 15 November 2010 |
| BRU | 1 | 2011 | 11 March 2011 | 24 May 2011 | NA |
| BRU | 1 | 2012 | 11 March 2012 | 30 May 2012 | 01 December 2012 |
| BRU | 1 | 2013 | 01 April 2013 | 11 July 2013 | 10 November 2013 |
| BRU | 1 | 2014 | 01 April 2014 | 19 June 2014 | 10 November 2014 |
| BRU | 2 | 2009 | 29 March 2009 | 18 June 2009 | 05 December 2009 |
| BRU | 2 | 2010 | 20 March 2010 | 07 June 2010 | 15 November 2010 |
| BRU | 2 | 2011 | 12 March 2011 | 24 May 2011 | NA |
| BRU | 2 | 2012 | 12 March 2012 | 30 May 2012 | 01 December 2012 |
| BRU | 2 | 2013 | 01 April 2013 | 11 July 2013 | 10 November 2013 |
| BRU | 2 | 2014 | 01 April 2014 | 19 June 2014 | 10 November 2014 |
| CHA | 1 | 2009 | 28 March 2009 | 15 July 2009 | 25 November 2009 |
| CHA | 1 | 2010 | 19 March 2010 | 09 June 2010 | 20 November 2010 |
| CHA | 1 | 2011 | 20 March 2011 | 28 June 2011 | NA |
| CHA | 1 | 2012 | 20 March 2012 | 26 June 2012 | 20 October 2012 |
| CHA | 1 | 2013 | 12 April 2013 | 18 July 2013 | NA |
| CHA | 2 | 2009 | 28 March 2009 | 15 July 2009 | 25 November 2009 |
| CHA | 2 | 2010 | 15 March 2010 | 09 June 2010 | 20 November 2010 |
| CHA | 2 | 2011 | 16 March 2011 | 28 June 2011 | NA |
| CHA | 2 | 2012 | 16 March 2012 | 26 June 2012 | 20 October 2012 |
| CHA | 2 | 2013 | 12 April 2013 | 18 July 2013 | NA |
| CHA | 3 | 2009 | 07 May 2009 | 16 July 2009 | 25 November 2009 |
| CHA | 3 | 2010 | 24 April 2010 | 09 June 2010 | 20 November 2010 |
| CHA | 3 | 2011 | 31 March 2011 | 28 June 2011 | NA |
| CHA | 3 | 2012 | 31 March 2012 | 26 June 2012 | 20 October 2012 |
| CHA | 3 | 2013 | 03 May 2013 | 18 July 2013 | NA |
| GAL | 1 | 2009 | 30 May 2009 | 06 August 2009 | 09 October 2009 |
| GAL | 1 | 2010 | 13 June 2010 | 19 July 2010 | 12 October 2010 |
| GAL | 1 | 2011 | 05 June 2011 | 18 July 2011 | 20 October 2011 |
| GAL | 1 | 2012 | 15 June 2012 | 24 July 2012 | 01 November 2012 |
| GAL | 1 | 2013 | 28 June 2013 | 15 August 2013 | 09 October 2013 |
| GAL | 1 | 2014 | 19 June 2014 | 06 August 2014 | 09 October 2014 |
| GAL | 2 | 2009 | 17 June 2009 | 06 August 2009 | 09 October 2009 |
| GAL | 2 | 2010 | 02 July 2010 | 19 July 2010 | 12 October 2010 |
| GAL | 2 | 2011 | 16 June 2011 | 18 July 2011 | 20 October 2011 |
| GAL | 2 | 2012 | 28 June 2012 | 24 July 2012 | 01 November 2012 |
| GAL | 2 | 2013 | 06 July 2013 | 15 August 2013 | 09 October 2013 |
| GAL | 2 | 2014 | 19 June 2014 | 06 August 2014 | 09 October 2014 |
| GAL | 3 | 2009 | 14 June 2009 | 06 August 2009 | 09 October 2009 |
| GAL | 3 | 2010 | 13 June 2010 | 19 July 2010 | 12 October 2010 |
| GAL | 3 | 2011 | 14 June 2011 | 18 July 2011 | 20 October 2011 |
| GAL | 3 | 2012 | 22 June 2012 | 24 July 2012 | 01 November 2012 |
| GAL | 3 | 2013 | 06 July 2013 | 15 August 2013 | 09 October 2013 |
| GAL | 3 | 2014 | 19 June 2014 | 06 August 2014 | 09 October 2014 |
| LAU | 1 | 2009 | 10 June 2009 | 05 August 2009 | 15 October 2009 |
| LAU | 1 | 2010 | 09 June 2010 | 18 July 2010 | 15 October 2010 |
| LAU | 1 | 2011 | 13 June 2011 | 10 August 2011 | 17 October 2011 |
| LAU | 1 | 2012 | 01 June 2012 | 23 July 2012 | NA |
| LAU | 1 | 2013 | 01 June 2013 | 15 August 2013 | NA |
| LAU | 1 | 2014 | 20 May 2014 | 06 August 2014 | NA |
| LAU | 2 | 2009 | 11 June 2009 | 05 August 2009 | 15 October 2009 |
| LAU | 2 | 2010 | 09 June 2010 | 18 July 2010 | 15 October 2010 |
| LAU | 2 | 2011 | 14 June 2011 | 10 August 2011 | 17 October 2011 |
| LAU | 2 | 2012 | 31 May 2012 | 23 July 2012 | NA |
| LAU | 2 | 2013 | 31 May 2013 | 15 August 2013 | NA |
| LAU | 2 | 2014 | 20 May 2014 | 06 August 2014 | NA |
| LAU | 3 | 2009 | 31 May 2009 | 05 August 2009 | 15 October 2009 |
| LAU | 3 | 2010 | 11 June 2010 | 18 July 2010 | 15 October 2010 |
| LAU | 3 | 2011 | 31 May 2011 | 19 July 2011 | 17 October 2011 |
| LAU | 3 | 2012 | 01 June 2012 | 23 July 2012 | NA |
| LAU | 3 | 2013 | 01 June 2013 | 15 August 2013 | NA |
| LAU | 3 | 2014 | 20 May 2014 | 06 August 2014 | NA |
| PIC | 1 | 2009 | 16 May 2009 | 11 August 2009 | 01 November 2009 |
| PIC | 1 | 2010 | 23 April 2010 | 03 August 2010 | 20 October 2010 |
| PIC | 1 | 2011 | 04 April 2011 | 10 August 2011 | 20 October 2011 |
| PIC | 1 | 2012 | 09 May 2012 | 24 July 2012 | 25 October 2012 |
| PIC | 1 | 2013 | 12 June 2013 | 16 August 2013 | 10 October 2013 |
| PIC | 1 | 2014 | 07 June 2014 | 07 August 2014 | 10 October 2014 |
| PIC | 2 | 2009 | 18 June 2009 | 11 August 2009 | 01 November 2009 |
| PIC | 2 | 2010 | 11 June 2010 | 03 August 2010 | 20 October 2010 |
| PIC | 2 | 2011 | 23 May 2011 | 10 August 2011 | 20 October 2011 |
| PIC | 2 | 2012 | 17 June 2012 | 24 July 2012 | 25 October 2012 |
| PIC | 2 | 2013 | 12 June 2013 | 16 August 2013 | 10 October 2013 |
| PIC | 2 | 2014 | 21 June 2014 | 07 August 2014 | 10 October 2014 |
| PIC | 3 | 2009 | 05 May 2009 | 11 August 2009 | 01 November 2009 |
| PIC | 3 | 2010 | 23 May 2010 | 03 August 2010 | 20 October 2010 |
| PIC | 3 | 2011 | 01 April 2011 | 10 August 2011 | 20 October 2011 |
| PIC | 3 | 2012 | 09 May 2012 | 24 July 2012 | 25 October 2012 |
| PIC | 3 | 2013 | 12 June 2013 | 16 August 2013 | 10 October 2013 |
| PIC | 3 | 2014 | 07 June 2014 | 07 August 2014 | 10 October 2014 |
| PIC | 4 | 2009 | 15 May 2009 | 11 August 2009 | 01 November 2009 |
| PIC | 4 | 2010 | 23 May 2010 | 03 August 2010 | 20 October 2010 |
| PIC | 4 | 2011 | 04 April 2011 | 10 August 2011 | 20 October 2011 |
| PIC | 4 | 2012 | 10 May 2012 | 24 July 2012 | 25 October 2012 |
| PIC | 4 | 2013 | 12 June 2013 | 16 August 2013 | 10 October 2013 |
| PIC | 4 | 2014 | 07 June 2014 | 07 August 2014 | 10 October 2014 |
| VIL | 1 | 2009 | 03 May 2009 | 30 July 2009 | 01 December 2009 |
| VIL | 1 | 2010 | 28 April 2010 | 13 July 2010 | 01 November 2010 |
| VIL | 1 | 2011 | 06 April 2011 | 15 July 2011 | 01 December 2011 |
| VIL | 1 | 2012 | 26 March 2012 | 09 July 2012 | 01 December 2012 |
| VIL | 1 | 2013 | 23 April 2013 | 22 July 2013 | 30 October 2013 |
| VIL | 1 | 2014 | 15 April 2014 | 18 July 2014 | 30 October 2014 |
| VIL | 2 | 2009 | 09 May 2009 | 29 July 2009 | 01 December 2009 |
| VIL | 2 | 2010 | 28 April 2010 | 13 July 2010 | 15 October 2010 |
| VIL | 2 | 2011 | 09 April 2011 | 15 July 2011 | 01 December 2011 |
| VIL | 2 | 2012 | 24 March 2012 | 09 July 2012 | 01 December 2012 |
| VIL | 2 | 2013 | 23 April 2013 | 22 July 2013 | 30 October 2013 |
| VIL | 2 | 2014 | 15 April 2014 | 18 July 2014 | 30 October 2014 |
| VIL | 3 | 2009 | 06 May 2009 | 29 July 2009 | 01 December 2009 |
| VIL | 3 | 2010 | 27 April 2010 | 13 July 2010 | 15 November 2010 |
| VIL | 3 | 2011 | 09 April 2011 | 15 July 2011 | 01 December 2011 |
| VIL | 3 | 2012 | 25 March 2012 | 09 July 2012 | 01 December 2012 |
| VIL | 3 | 2013 | 23 April 2013 | 22 July 2013 | 30 October 2013 |
| VIL | 3 | 2014 | 15 April 2014 | 18 July 2014 | 30 October 2014 |

**Table S3.** Pearson’s linear correlation coefficients between early season and late season thermic variables. Correlations higher than 0.7 are in bold.

|  | T.min.1 | T.mean.1 | T.max.1 | DD.1 | T.range.1 | Frost.1 | Frost.e.1 |
| --- | --- | --- | --- | --- | --- | --- | --- |
| T.min.1 | 1.00 | 0.67 | -0.02 | 0.09 | -0.49 | -0.66 | -0.67 |
| T.mean.1 | 0.67 | 1.00 | **0.71** | 0.45 | 0.30 | -0.37 | -0.36 |
| T.max.1 | -0.02 | **0.71** | 1.00 | 0.59 | **0.88** | 0.08 | 0.10 |
| DD.1 | 0.09 | 0.45 | 0.59 | 1.00 | 0.47 | 0.10 | 0.10 |
| T.range.1 | -0.49 | 0.30 | **0.88** | 0.47 | 1.00 | 0.39 | 0.41 |
| Frost.1 | -0.66 | -0.37 | 0.08 | 0.10 | 0.39 | 1.00 | **0.99** |
| Frost.e.1 | -0.67 | -0.36 | 0.10 | 0.10 | 0.41 | **0.99** | 1.00 |

|  | T.min.2 | T.mean.2 | T.max.2 | DD.2 | T.range.2 | Frost.2 | Frost.e.2 |
| --- | --- | --- | --- | --- | --- | --- | --- |
| T.min.2 | 1.00 | **0.79** | 0.12 | **0.77** | -0.47 | -0.55 | -0.61 |
| T.mean.2 | **0.79** | 1.00 | 0.69 | **0.89** | 0.16 | -0.27 | -0.31 |
| T.max.2 | 0.12 | 0.69 | 1.00 | 0.52 | 0.82 | 0.14 | 0.15 |
| DD.2 | **0.77** | **0.89** | 0.52 | 1.00 | 0.02 | -0.23 | -0.29 |
| T.range.2 | -0.47 | 0.16 | 0.82 | 0.02 | 1.00 | 0.44 | 0.49 |
| Frost.2 | -0.55 | -0.27 | 0.14 | -0.23 | 0.44 | 1.00 | **0.98** |
| Frost.e.2 | -0.61 | -0.31 | 0.15 | -0.29 | 0.49 | **0.98** | 1.00 |

**Figure S1.**
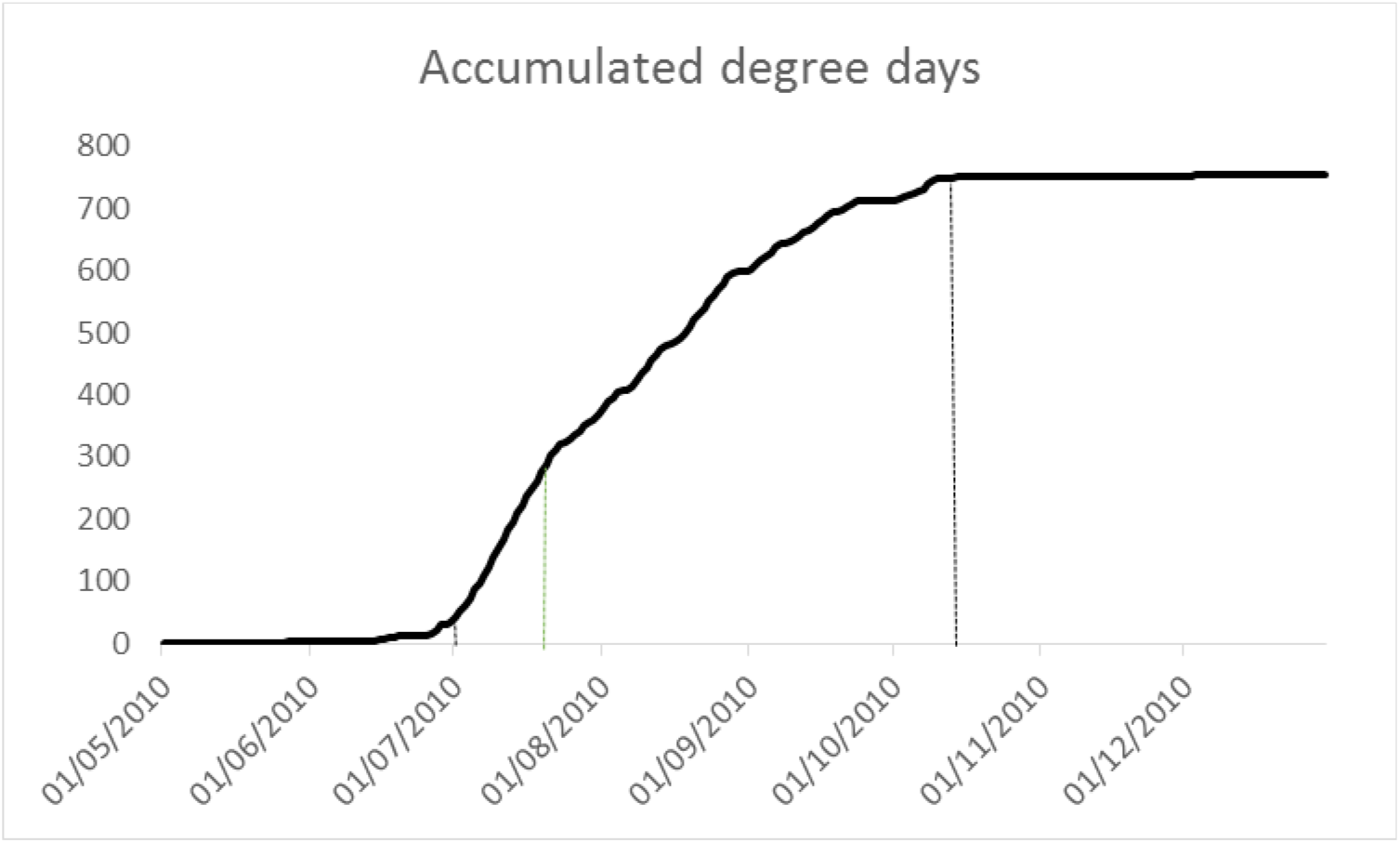
Example of an accumulated degree days curve used to define the limits of the vegetative season (vertical black lines). The date of census is also showed (green vertical line)

